# Glucocorticoids regulate the human non-coding genome

**DOI:** 10.64898/2025.12.18.695158

**Authors:** Thai Tran, Robert Kwiat, Qilin Cao, Skyler Kuhn, Katherine N. Howe, Manasi Gadkari, Luis M. Franco

## Abstract

Glucocorticoids (GCs) are widely used anti-inflammatory and immunosuppressive agents known to induce dramatic changes in gene expression, yet their effects on non-coding RNAs such as long non-coding RNAs (lncRNAs) or microRNAs (miRNAs) remain poorly characterized. We present the first comprehensive and systematic analysis of GC regulation of the non-coding genome across nine human primary cell types from both hematopoietic and non-hematopoietic lineages. In vitro GC treatment was applied to each cell type, with transcriptomic profiling (total RNA-seq and small RNA-seq) at 2 and 6 hours post-treatment. We identified over 2,000 GC-responsive non-coding transcripts, including 654 annotated lncRNAs, 1,376 novel lncRNAs, and 39 miRNAs. The non-coding RNA response to GCs was highly cell type-dependent: 80% of GC-responsive lncRNAs and 97% of miRNAs were unique to a single cell type. Hematopoietic cells exhibited a greater magnitude of lncRNA induction than non-hematopoietic cells. Notably, dozens of facultative lncRNAs (undetectable at baseline) were induced de novo by GC. GC-responsive transcripts spanned diverse lncRNA classes, with enrichment of host lncRNAs. By contrast, only a limited number of miRNAs were GC-responsive. GC regulation of transcript abundance for miRNAs and their host lncRNAs appears to be independent. Our results, which are accessible through an interactive web application, establish a framework for studying how non-coding transcription contributes to physiological and clinical heterogeneity in GC responses. The newly identified GC-responsive non-coding transcripts could represent biomarkers of GC exposure, determinants of GC sensitivity or resistance, or candidate regulators of tissue-specific GC effects.

## Introduction

Glucocorticoids (GCs) are the cornerstone of immunosuppressive and anti-inflammatory therapies in humans. Despite their widespread use in clinical medicine, there are still substantial gaps in our understanding of the mechanisms by which GCs regulate immunity (Cain and Cidlowski, 2017, Bhatt and Franco, 2025). The glucocorticoid receptor (GR; UniProt: P04150), encoded by the gene *NR3C1*, is a transcription factor of the steroid/thyroid hormone nuclear receptor superfamily. The ligand-bound GR can dimerize and directly bind DNA at specific recognition sequences known as glucocorticoid response elements (GREs), increasing transcription rates. It can also be recruited to specific sites in the genome via protein-protein interactions with other DNA-bound transcription factors (Sacta et al., 2016). Monomeric GR has been proposed to bind DNA at a distinct set of recognition sequences known as negative GREs, decreasing transcription rates (Hudson et al., 2013, Surjit et al., 2011). Other molecular mechanisms of GC action have been described, including direct biochemical effects on the tricarboxylic acid cycle and its metabolites (Auger et al., 2024, Stifel et al., 2022, Franco, 2024), alterations of ion transport across membranes (Buttgereit et al., 1993, Schmid et al., 2000), and interactions with proposed membrane-bound forms of GR (Bartholome et al., 2004, Gametchu, 1987, Gametchu et al., 1993). Although the molecular mechanisms are diverse, a consistent outcome of GC treatment is a substantial reprogramming of a cell’s transcriptional state (Galon et al., 2002, Olnes et al., 2016, Franco et al., 2019). This strong transcriptional response is highly cell type-dependent (Franco et al., 2019), at least in part due to differences in chromatin accessibility and expression of GR co-factors (John et al., 2011, Reddy et al., 2012, Grontved et al., 2013).

While the existing evidence indicates that transcriptional reprogramming is the main molecular mechanism of GC action, studies to date have almost exclusively focused on the response of the protein-coding elements of the genome. However, a limited but growing body of experimental evidence indicates that GC treatment leads to changes in the abundance of human non-coding transcripts, with potential functional implications. For example, the micro-RNA miR-155, which was initially found to be reduced by GC treatment in mouse peritoneal macrophages and liver samples in lipopolysaccharide (LPS)-induced sepsis (Wang et al., 2013, Zheng et al., 2012), was later found to also be reduced after GC treatment in human primary CD4+ T cells stimulated with dust mite extract (Daniel et al., 2018). Given that miR-155 has been found to drive pro-inflammatory cytokine production, while suppressing negative regulators of inflammation (Kurowska-Stolarska et al., 2011, Cardoso et al., 2012), its suppression has been postulated as a possible contributor to the anti-inflammatory effects of GCs. miRNA-155 is also a good example of the cell-type dependence of GC responses, as it has found to be induced rather than repressed in 3T3-L1 cells upon GC exposure (Peshdary and Atlas, 2018). Similarly, the microRNA miR-511 was initially found to be GC-induced in mouse liver and spleen, leading to reduced expression of a target gene encoding the TNF receptor TNFR1 (Puimège et al., 2015). The mature form miR-511-5p was later documented to also be induced by GCs in human circulating monocytes, where the LPS receptor TLR4 appears to be a key target (Curtale et al., 2017). The microRNA miR-98 has also been shown to be induced by GCs in human primary T lymphocytes, leading to reduced abundance of predicted targets including transcripts for interleukin-13 and multiple TNF receptors (Davis et al., 2013). The microRNA miR-124 has also been documented to be GC-responsive in human primary T cells, resulting in decreased expression of the alpha isoform of the GR and potentially contributing to clinical GC resistance (Ledderose et al., 2012). Fewer studies have looked at the effects of GCs on human long non-coding RNAs (lnc-RNAs), but there is experimental evidence of GC induction of the lncRNA *DANCR* in human adipose-derived mesenchymal stem cells (Yan et al., 2025) and of GC repression of a non-coding transcript variant of the gene *TCF7L2* in human induced pluripotent stem cell-derived astrocytes (Liu et al., 2021). Although not yet documented to be GC targets in human primary cells, additional non-coding transcripts have been found to be GC responsive in cancer cell lines or animal models (Kim et al., 2020, Rapicavoli et al., 2013, Casciaro et al., 2023, Liu et al., 2018, Mirzadeh Azad et al., 2019, Zhu et al., 2010).

While these studies provide clear evidence of a relationship between GCs and non-coding elements of the genome, a comprehensive and systematic characterization of the effects of GCs on the human non-coding genome is not yet available, as has been pointed out in recent reviews (Syed et al., 2020, Pierouli et al., 2022). To address this substantial knowledge gap, we have studied the transcriptional response of non-coding elements of the genome to GCs in nine human primary cell types.

## Results

### A systematic assessment of the response to glucocorticoids by the human non-coding genome

We systematically assessed the response of non-coding elements of the genome to GCs in nine human primary cell types obtained from healthy donors: B cells, CD4+ T cells, endothelial cells, fibroblasts, monocytes, myoblasts, neutrophils, osteoblasts, and preadipocytes (**Figure 1**). Each cell type was treated in vitro with methylprednisolone or vehicle and sampled at 2 and 6 hours. RNA was purified and separately processed for total RNA sequencing (n = 4) or small-RNA sequencing (n = 3). The total RNA sequencing (RNA-seq) dataset was used for analysis of long non-coding transcripts, and the small-RNA-seq dataset for analysis of microRNAs (miRNAs). To explore the possibility of GC-responsive novel (not previously annotated) non-coding transcripts, we performed de novo transcript assembly with the total RNA-seq alignments (**Figure S1A**) and analyzed the small-RNA-seq data with a program (Friedlander et al., 2008) that identifies known or novel miRNAs in next-generation sequencing data (**Figure S1B**). This analysis revealed 2,069 non-coding elements in the human genome that were GC-responsive in one or more cell types, at one or both time points: 654 GENCODE-annotated lncRNAs, 1,376 novel de novo assembled transcripts, 35 miRbase-annotated miRNAs, and 4 predicted novel miRNAs (**Figure 1**).

**Figure 1.**
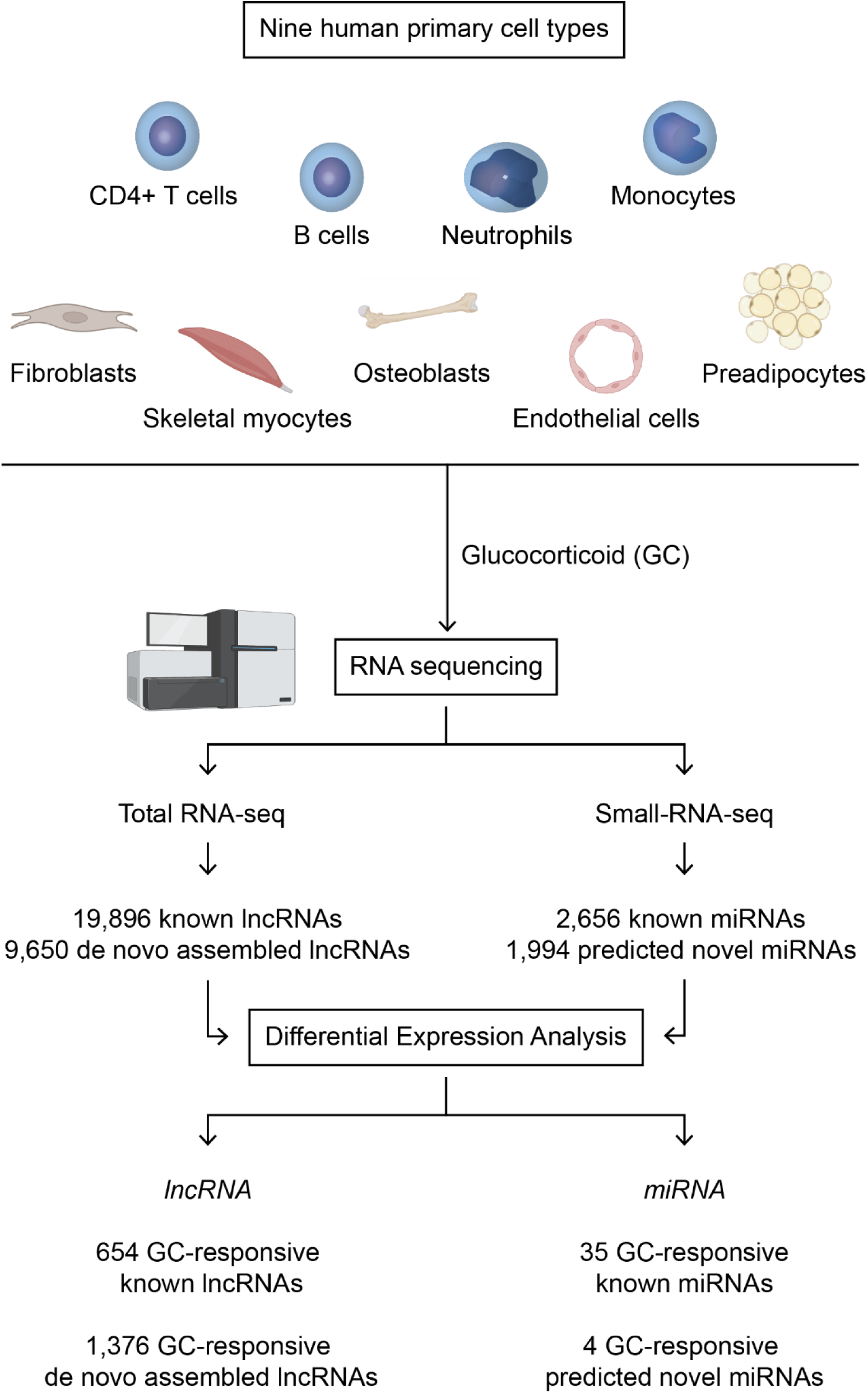
Experimental design. Human primary cells were obtained from healthy donors. The number of biological replicates (unrelated donors) for each cell type was 4 for total RNA sequencing and 3 for small-RNA sequencing. In vitro treatment was performed independently for each cell type and biological replicate with glucocorticoid (GC; methylprednisolone) or vehicle (ethanol). Cells were sampled 2 and 6 hours after treatment. Differential expression analysis involved GC versus vehicle contrasts at each time point.

### Glucocorticoids regulate the non-coding genome in a cell type-dependent manner

GCs regulate the non-coding human genome (**Figure 2**). The number of GC-responsive lncRNAs varies by cell type; this is true for GENCODE-annotated, as well as novel (de novo assembled), lncRNAs (**Figure 2A**). Overall, the transcriptional response to GCs by the human non-coding genome was stronger in hematopoietic than in non-hematopoietic cells (**Figures 2A and 2B**). As we had previously described for coding genes (Franco et al., 2019), neutrophils are the cell type most transcriptionally responsive to GCs at the level of lncRNAs, while endothelial cells are the least responsive (**Figure 2A**).

**Figure 2.**
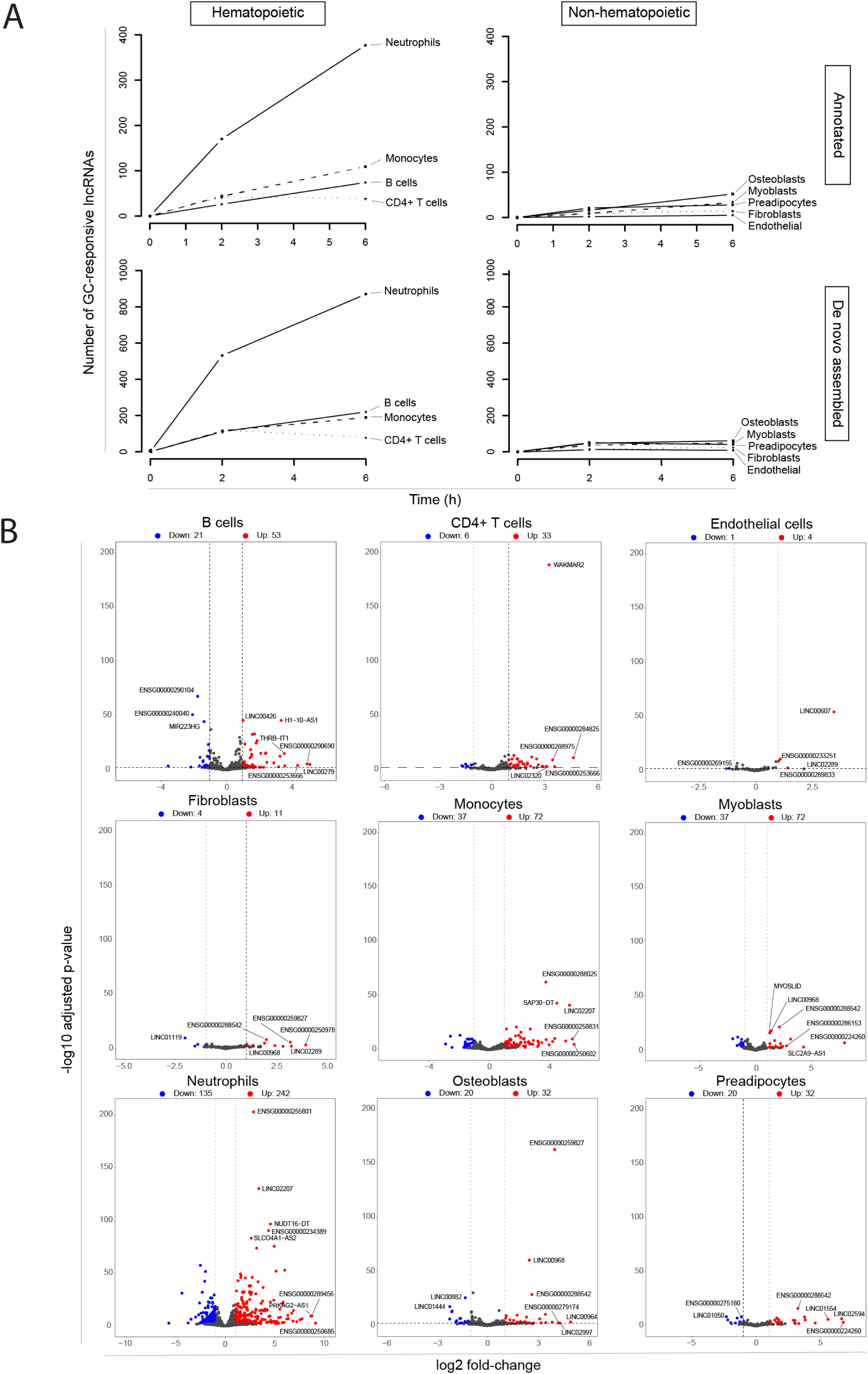
Glucocorticoids regulate the non-coding human genome. For each of nine primary human cell types, total RNA was purified 2 and 6 hours after in vitro administration of methylprednisolone (22.7 µM) or vehicle (ethanol 0.08%). Results of four biological replicates (unrelated healthy donors). RNA sequencing and differential expression analysis were performed, contrasting methylprednisolone-treated versus vehicle-treated cells at each time point. **(A)** Number of glucocorticoid-responsive long non-coding RNAs over time in hematopoietic or non-hematopoietic cell types. A glucocorticoid-responsive lncRNA is defined as one with an adjusted p-value for differential expression ≤ 0.01 and log2 fold-change > 1 or < −1. Results are shown separately for GENCODE-annotated lncRNAs (top), or de novo assembled lncRNAs (bottom). **(B)** Volcano plots of the response to glucocorticoids by cell type. Results are shown for GENCODE-annotated lncRNAs at the 6-hour time point. Long ncRNAs with significantly increased or reduced transcript abundance are highlighted in orange or blue, respectively. Significantly increased transcript abundance is defined as an adjusted p-value for differential expression ≤ 0.01 and log2 fold-change > 1. Significantly decreased transcript abundance is defined as an adjusted p-value for differential expression ≤ 0.01 and log2 fold-change < −1. Results for de novo annotated lncRNAs are shown in **Figure S2**.

Among GC-responsive lncRNAs, at the time points tested, there is a predominance of transcripts with increased abundance after GC administration in all cell types studied. This pattern was observed in both annotated (**Figure 2B**) and de novo assembled (**Figure S2**) lncRNAs.

Analysis of the specific lncRNA transcripts that are GC-responsive in each cell type also reveals a striking cell type dependence. Of the 564 annotated lncRNAs that are GC-responsive in one or more hematopoietic cell types, 476 (84.4%) are responsive in only one cell type (**Figure 3A**, top left). Of the 108 annotated lncRNAs that are GC-responsive in one or more non-hematopoietic cell types, 73 (67.6%) are responsive in only one cell type (**Figure 3A**, top right). Similarly, of the 1,291 de novo assembled lncRNAs that are GC-responsive in one or more hematopoietic cell types, 1,098 (85%) are responsive in only one cell type (**Figure 3A**, bottom left). Of the 137 de novo assembled lncRNAs that are GC-responsive in one or more non-hematopoietic cells, 77 (56.2%) are responsive in only one cell type (**Figure 3A**, bottom right).

**Figure 3.**
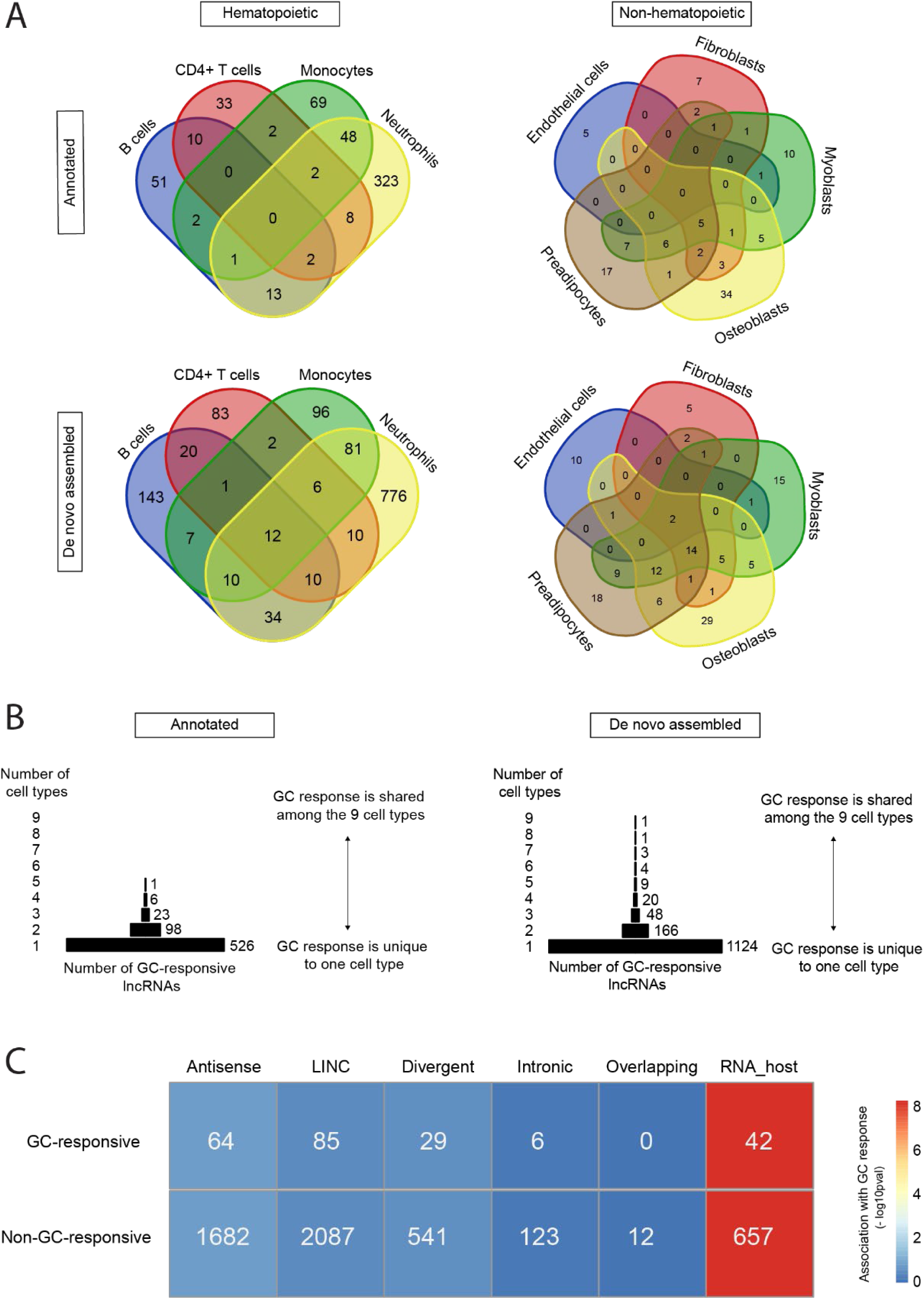
The response of human lncRNAs to GCs varies by cell type and by lncRNA subtype. **(A)** Venn diagrams of the cell type distribution of GC-responsive lncRNAs. **(B)** Pyramid plot of GC-responsive lncRNAs by the number of cell types in which they were found to be GC-responsive. In panels A and B, a GC-responsive lncRNA was defined as one with an adjusted p-value for differential expression ≤ 0.01, and a log2 fold-change ≥ 1 or ≤ −1, at the 2-hour or 6-hour time point. The lists are non-redundant: if a lncRNA is GC-responsive at 2 and 6 hours, it is counted once. **(C)** GC response by different lncRNA subtypes. Association was tested by Fisher’s exact test. The heatmap displays the strength of association as the

The cell type-dependence of the response to GCs by human non-coding transcripts appears to be more extreme than what we had previously described for coding transcripts (Franco et al., 2019). Of the 654 GC-responsive lncRNAs that we have identified, none are GC-responsive in more than five cell types and 526 (80%) are GC-responsive in only one of the nine cell types studied (**Figure 3B**). Similarly, among the 1,176 GC-responsive de novo assembled lncRNAs, only one transcript is GC-responsive in all nine cell types, and 1,124 (81.7%) are GC-responsive in only one of the nine cell types studied (**Figure 3B**).

### The ‘host RNA’ category of lncRNAs is associated with glucocorticoid responsiveness

We then assessed whether specific subtypes of lncRNAs are more likely than others to be GC-responsive. The HUGO Gene Nomenclature Committee has recently published an updated standardized nomenclature for lncRNAs (Seal et al., 2023), which classifies them based on their genomic location and direction with respect to other genes (Seal et al., 2020). Based on this, we considered 6 categories of lncRNAs: antisense lncRNAs, long intergenic non-coding RNAs (LINC), divergent lncRNAs, intronic lncRNAs, overlapping lncRNAs, and lncRNAs that serve as host genes for miRNAs or small nucleolar ncRNAs (snoRNAs). It’s worth noting that, although classification efforts have advanced substantially in recent years, the available annotation of lncRNAs for studies like this remains incomplete. Of the 19,928 records in the GENCODE v43 annotation file for lncRNAs, 5,328 (26.7%) are annotated further to indicate that they belong to one or more of the six categories above. Of the 654 GC-responsive annotated lncRNAs, 226 are annotated further. A breakdown of the number GC responsive transcripts by lncRNAs category is displayed in **Figure 3C**. The ‘host RNA’ category was significantly associated with GC response (Fisher’s exact test p = 4.9 × 10^-9^). We found no statistical evidence of an association between GC response and any of the other lncRNA categories.

### Glucocorticoid treatment uncovers facultative, stimulus-dependent lncRNAs

Given the extensive effects of GCs on transcriptional activity, we wondered if there could be regions of the human genome that are not expressed at baseline but transcribed upon GC treatment. Such transcripts could represent facultative lncRNAs which are unlikely to have been discovered previously, given that their expression is stimulus-dependent. Across the 9 cell types and 2 time points, we identified 49 unique GC-responsive de novo assembled lncRNAs with unnormalized read counts of zero in the vehicle condition. Five cell types (monocytes, myoblasts, neutrophils, osteoblasts, and preadipocytes) had at least one candidate stimulus-dependent lncRNA. Consistent with their strong GC responsiveness, neutrophils are the cell type with the highest number of candidate stimulus-dependent lncRNAs. **Figure 4** provides two examples of facultative lncRNAs uncovered by GC treatment.

**Figure 4.**
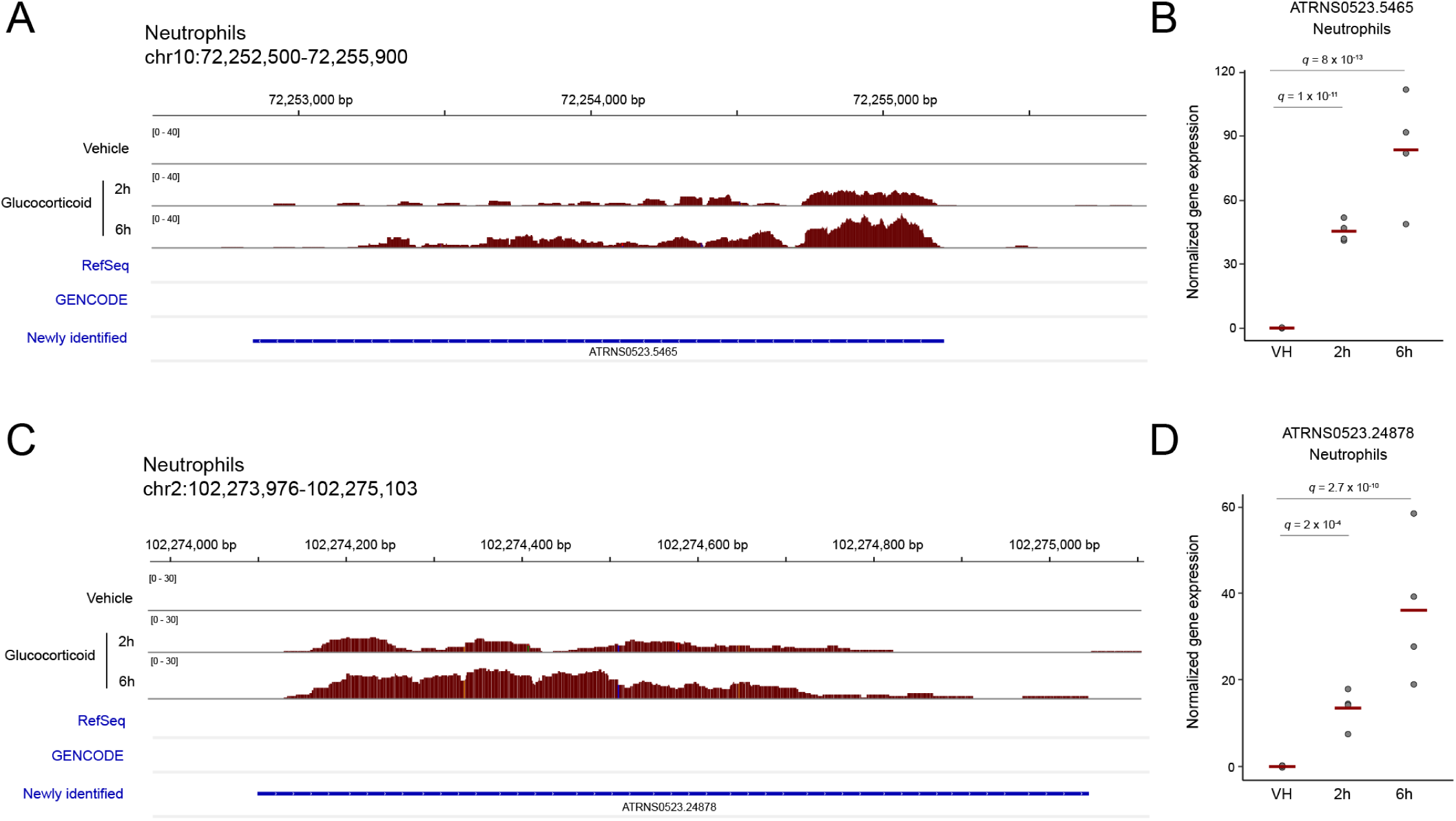
GC treatment uncovers facultative, stimulus-dependent lncRNAs. **(A)** An unannotated region of human chromosome 10 with a newly identified lncRNA that shows no evidence of expression at baseline and transcriptional induction in response to GC treatment in primary neutrophils. **(B)** Normalized read count and adjusted p-value (q) for differential expression, for the newly described lncRNA displayed in panel A. **(C)** An unannotated region of human chromosome 2 with a newly identified lncRNA that shows no evidence of expression at baseline and transcriptional induction in response to GC treatment in primary neutrophils. **(D)** Normalized read count and adjusted p-value (q) for differential expression, for the newly described lncRNA displayed in panel C. Panels A and C were generated with the Integrative Genomics Viewer (IGV).

### The response of human miRNAs to glucocorticoids is limited and cell type-dependent

Our small-RNA-seq analysis identified 1,729 known (miRBase-annotated) miRNAs and 1,994 predicted novel (not miRBase-annotated) miRNAs with evidence of expression in at least one sample across the 9 cell types. The overall pattern of miRNA expression is characteristic of different cell types. Principal component analysis (PCA) of the known-miRNA expression values results in clusters corresponding to the ontogenetic origin of the 9 human primary cell types studied, with the first component separating hematopoietic from non-hematopoietic cells, defined clusters for lymphoid cells, and a separation of endothelial cells from other non-hematopoietic cells (**Figure 5A**). The same pattern is seen with the expression values from the predicted novel miRNAs (**Figure 5B**), indicating that this set of miRNAs is likely to reflect biologically relevant molecules and not random noise in the sequencing data.

**Figure 5.**
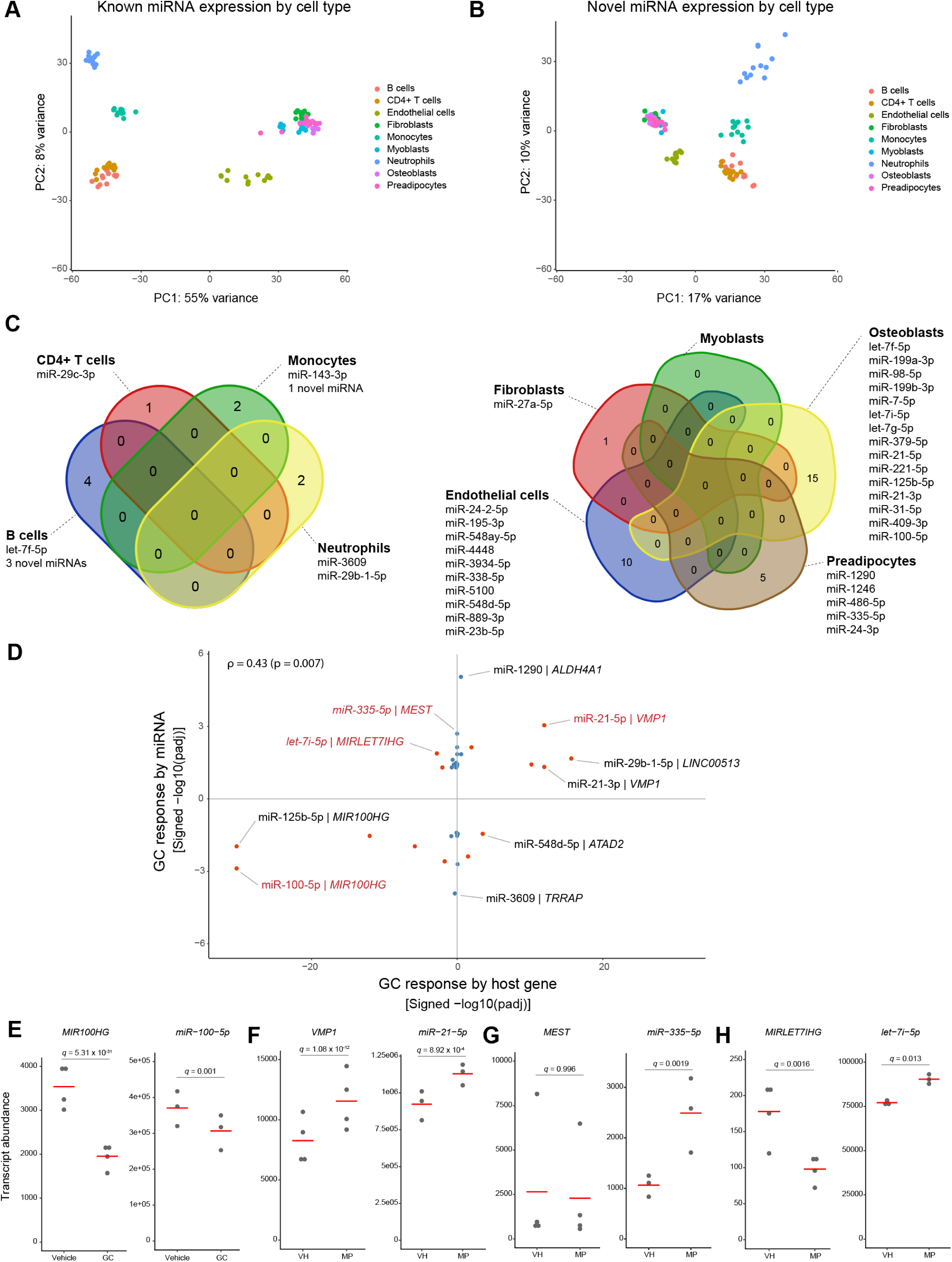
The response of human miRNAs to GCs is limited and cell type-dependent. **(A)** PCA of miRNA expression for known (miRbase-annotated) miRNAs in 9 human primary cell types. **(B)** PCA of miRNA expression for predicted novel miRNAs in 9 human primary cell types. **(C)** GC-responsive miRNAs in 4 human hematopoietic cell types (left) and 5 human non-hematopoietic cell types (right). **(D)** GC response by intragenic miRNAs and their host genes. Each dot represents one miRNA/host-gene pair. Pairs in which both miRNA and host gene were GC-responsive are marked in red. Rho = Pearson correlation coefficient. **(E-F)** Examples of miRNA/host-gene pairs in which the GC response is congruent. **(G)** Example of a miRNA/host-gene pair in which GC regulation is independent (miRNA is GC-responsive, while the host gene is not). **(H)** Example of a miRNA/host-gene pair in which the GC response is opposite.

Of the 1,729 known miRNAs identified, only 10 were found to be GC-responsive in one or more cell types at either time point, when applying the same adjusted p-value threshold of p ≤ 0.01 that we employed in the analysis of the lncRNA data. Those are MIR29C (hsa-mir-29c-3p mature miRNA) in CD4+ T cells; MIR143 (hsa-mir-143-3p) in monocytes; MIR3609 (hsa-mir-3609) in neutrophils; MIR27A (hsa-mir-27a-5p) in fibroblasts; MIRLET7F (hsa-let-7f-5p), MIR100 (hsa-miR-100-5p) and MIR21 (hsa-miR-21-5p) in osteoblasts; and MIR1290 (hsa-miR-1290), MIR1246 (hsa-miR-1246) and MIR335 (hsa-miR-335-5p) in preadipocytes. At that threshold, none of the predicted novel miRNAs can be classified as GC-responsive. Applying a less stringent threshold of p ≤ 0.05, 39 miRNAs (35 known and 4 predicted) are found to be GC-responsive in one or more cell types (**Figure 5C and Table S1**). The cell type-dependence of the transcriptional response to GCs is even more striking for miRNAs than it is for long non-coding or coding transcripts: only 1 of the 39 GC-responsive miRNAs, MIRLET7F (hsa-let-7f-5p), is GC-responsive in more than one cell type (B cells and osteoblasts), while the others are cell type-specific (**Figure 5C**).

We then assessed whether the GC-responsive miRNAs were primarily intragenic (encoded by a sequence that is within a coding or non-coding long-RNA host) or intergenic. A majority of known mammalian miRNAs are intragenic, with a current estimate around 69% (Rodriguez et al., 2004). Of the 39 GC-responsive miRNAs, 34 (87.2%) are intragenic, 4 (10.2%) are intergenic, and 1 (2.6%) is encoded by both intra- and intergenic regions. Of the 34 GC-responsive miRNAs that are intragenic, 19 (55.9%) have lncRNA host genes, 14 (41.2%) have coding host genes, and 1 (2.9%) is encoded by a coding gene and a non-coding host gene (**Table S1**).

Finally, we studied the relationship between the GC response of intragenic miRNAs and that of their host genes. Among the GC-responsive mature miRNAs that are encoded by at least one intragenic region, there is a weak correlation (Pearson r = 0.43, p = 0.007) between the GC response of the host gene and that of the mature miRNA (**Figure 5D**). For some miRNAs, there is clear evidence of co-regulation, whereas for most miRNAs, the response is not accompanied by a significant transcriptional response to GCs by the host gene, indicating independent regulation. The GC response was congruent (increased or decreased transcript abundance in both miRNA and host gene in the given cell type and time point) in 10 miRNA/host-gene pairs (26%); the response was opposite (increased expression in miRNA and decreased expression in host gene, or vice-versa) in 4 (10.5%), and only the miRNA was GC-responsive in the remaining 24 (63%) (**Figure 5D**). Specific examples of congruent, opposite, and independent regulation by GCs of miRNAs and their host genes, are shown in **Figure 5 E-H**.

### An interactive tool for exploring the response of human non-coding genes to glucocorticoids

For this comprehensive and systematic assessment of the effects of GCs on the non-coding human genome to be of maximum use to the research community, we have developed a web application for interactively querying the study’s results (<u>Interactive</u> Figure 1). Users can select a cell type and generate displays of the genome-wide response of non-coding transcripts to GCs, with user-selected cut-off values for differential expression p-value and log fold-change. Users can also highlight specific non-coding genes of interest or zoom into specific segments of the genome and display the relevant data for GC-responsive non-coding transcripts (known or newly identified) in tabular format. It is our hope that this tool will allow other investigators to initiate their own explorations of the transcriptional response of the non-coding genome to GCs and be a starting point for studies of the potential functional consequences of this response in different biological contexts.

## Discussion

Despite their clinical and physiological importance, substantial gaps exist in our understanding of the molecular mechanisms by which GCs regulate biological functions. Much of the existing work has involved GC effects on coding elements of the genome and has been performed on cancer cell lines, which differ biologically from human primary cells. GC effects on non-coding RNAs have been reported sporadically, but never comprehensively or across human primary cell types. This work directly addresses that knowledge gap, providing the first systematic assessment of GC-responsive non-coding transcripts (lncRNAs and mi-RNAs) across nine hematopoietic and non-hematopoietic human primary cell types.

Acute GC exposure induces extensive and highly cell type-dependent regulation of lncRNAs. We identified over 2,000 GC-responsive non-coding elements: 654 previously annotated lncRNAs and 1,376 novel (de novo assembled) lncRNAs. Human hematopoietic cells show a substantially stronger lncRNA response than non-hematopoietic cells, paralleling prior findings for coding genes. The cell specificity of the lncRNA response to GCs is remarkable, with 80% of GC-responsive lncRNAs being responsive in only one of the nine cell types studied. This cell-type specificity is even stronger than what we had previously described for coding elements of the genome (Franco et al., 2019) and it suggests that chromatin context and lineage-defining transcription factors likely constrain GC-mediated lncRNA induction.

We found a predominance of GC-induced lncRNAs across all cell types. Interestingly, this includes novel, facultative, treatment-dependent lncRNAs: we identified 49 novel lncRNAs with no detectable baseline transcription across cell types, which appear only upon acute GC treatment. This indicates that portions of the human non-coding genome are GC-inducible and therefore invisible in standard steady-state transcriptomics experiments. This finding parallels that of GC-induced enhancer RNAs (eRNAs) in murine macrophages (Greulich et al., 2021) and human cell lines (Mostafa et al., 2021), although our study did not specifically document eRNAs.

Among lncRNA subtypes, there is clear evidence of a GC response among intronic, intergenic, antisense, divergent, and host lncRNAs, although only host lncRNAs are selectively enriched. Because host RNAs can give rise to miRNAs or snoRNAs, it is possible that GCs can simultaneously coordinate the production of both lncRNAs and small RNAs embedded within them.

Although nearly all of the previously documented effects of GCs on the non-coding genome involved miRNAs, our results indicate that human lncRNAs are far more extensively regulated by GCs than miRNAs. Our unbiased analysis of small RNAs revealed only a small number of GC-responsive miRNAs in the primary cells studied: Of the 1,729 known miRNAs identified, 10 were found to be GC-responsive in one or more cell types when applying the same adjusted p-value threshold of p ≤ 0.01 that we employed in the analysis of the lncRNA data. That number increased to 39 (35 known and 4 novel miRNAs) with a relaxed threshold of 0.05. Again, the response was highly cell type-dependent, with 38 of 39 GC-responsive miRNAs regulated in a single cell type. Thus, miRNA regulation by GCs could depend on chromatin context, lineage-specific transcription factors, or cell-selective post-translational processing programs. Of the GC-responsive miRNAs identified, 87% are intragenic and, of those, more than half have a lncRNA host gene. However, the GC responsiveness of miRNAs correlates only weakly with that of their host genes (r = 0.43). We found clear evidence of congruent, independent, and even opposite directions of GC response by miRNAs and their host genes, suggesting independent control of miRNA expression, maturation, stability, or promoter usage.

Three limitations of this work are important to discuss. First, while it greatly broadens our understanding of the human non-coding transcripts that respond to GCs, substantial additional work will be required to establish which of these transcripts have functional consequences in individual cell types upon GC treatment. Second, given the known context-dependence of the cellular response to GCs, and the fact that different platforms for measurement of RNA abundance have varying detection characteristics, our results should not be misunderstood as providing evidence for or against a potential GC response by a non-coding gene in another species, cell type, biological condition, or measurement platform. If a given non-coding RNA has been found to be GC-responsive under different conditions, its absence from our results should not be misinterpreted as evidence against it as a GC target under those conditions. Third, our experimental design only examined acute GC exposure in vitro, and chronic GC effects could differ.

In summary, while the study of protein-coding genes has dominated the GC literature, our findings indicate that GCs strongly regulate the non-coding human genome in a cell type-dependent manner. Our results, which we have aimed to make publicly and readily available through an interactive web application, establish a framework for studying how non-coding transcription contributes to physiological and clinical heterogeneity in GC responses. It is our hope that these results will help conceptually shift the field toward a more integrated regulatory view where non-coding transcription is not peripheral but foundational. Because transcriptional responses to GCs underlie many of their known immunosuppressive effects and side effects, the newly identified GC-responsive non-coding transcripts could represent biomarkers of GC exposure, determinants of GC sensitivity or resistance, or candidate regulators of tissue-specific GC effects.

## Methods

### Cell purification and cell culture

Human peripheral blood hematopoietic cells were obtained from the Department of Transfusion Medicine at the National Institutes of Health (NIH) Clinical Center, under NIH study 99-CC-0168. PBMCs were isolated from a leukapheresis sample for biological replicate 1 and from peripheral blood collected in Vacutainer EDTA tubes (Becton Dickinson; cat. no. 366643) for all other replicates. Mononuclear cell subsets were isolated from PBMCs in SepMate tubes (STEMCELL Technologies; cat. no. 15460) by gradient centrifugation with Ficoll-Paque PLUS (GE Healthcare Life Sciences; cat. no. 17-1440-03) followed by immunomagnetic enrichment for the specific cell subset with EasySep Human cell enrichment kits (STEMCELL Technologies). B lymphocytes and CD4+ T lymphocytes were isolated by negative selection (STEMCELL Technologies; cat. nos. 19054 and 19052, respectively). Monocytes were isolated by positive selection (STEMCELL Technologies; cat. nos. 18058 or 17858), to ensure inclusion of the CD14+/CD16+ fraction. Neutrophils were isolated directly from whole blood by negative selection (STEMCELL Technologies; cat. no. 19666). Immediately after isolation, mononuclear cells were incubated overnight in 12-well plates (Corning; cat. no. 3512), in Advanced RPMI 1640 medium (Thermo Fisher Scientific; cat. no. 12633-012) with 1 x L-glutamine, 10 mM HEPES, and 10% autologous serum, at 37°C and 5% CO2. Neutrophils were incubated for 4 h in 12-well plates (Corning; cat. no. 3512) pre-coated with 20% autologous plasma, in RPMI 1640 medium without phenol red (Thermo Fisher Scientific; cat. no. 11835030) with 10 mM HEPES, at 37°C and 5% CO2.

Human primary non-hematopoietic cells were isolated from four unrelated healthy donors and cultured in 12-well plates in respective medium without antibiotics or GCs, at 37°C, 5% CO2. Human primary skeletal muscle myoblasts (Lonza; cat. no. CC-2580; lot nos. 0000419228, 0000650386, 0000657512, and 0000583849) were cultured in SkGM-2 medium (Lonza; cat. no. CC-3245). Human primary subcutaneous preadipocytes (Lonza; cat. no. PT-5020, lot nos. 0000399826, 0000629514, and 0000645827; Cell Applications, Inc.; cat. no. 802s-05a, lot no. 1687) were cultured in PGM-2 medium (Lonza; cat. no. PT-8002). Human dermal microvascular endothelial cells (Lonza; cat. no. CC-2543; lot nos. 0000442486, 0000577921, 0000550175, and 0000664503) were cultured in EGM-2MV medium (Lonza; cat. no. CC-3202). Human primary dermal fibroblasts (Lonza; cat. no. CC-2511; lot nos. 0000409270, 0000509796, 0000540991, and 0000545147) were cultured in FGM-CD medium (Lonza; cat. no. 00199041) or HyClone Minimum Essential Medium (Thermo Fisher Scientific; cat. no. SH30265.FS) with 10% FBS. Human primary osteoblasts (Lonza; cat. no. CC-2538, lot nos. 0000435102, 0000336963 and 0000426160; iXCells Biotechnologies; cat. no. 10HU-179, lot no. 200211) were cultured in OGM medium (Lonza; cat. no. CC-3207). A growth curve was generated to ensure GC treatment was performed in the early plateau phase of growth and cells were in passage 3 to 5 at the time of treatment.

### In vitro glucocorticoid treatment

For each cell type and biological replicate, in vitro treatment was performed independently with methylprednisolone 22.7 µM (Sigma; cat. no. M0639) or vehicle (ethanol, 0.08%) and sampled at 2 and 6 h after the stimulus. Immediately after harvesting, cells were centrifuged at 500 g for 5 minutes. The cell pellet was resuspended in 500 μl of TRIzol Reagent (Thermo Fisher Scientific; cat. no. 15596018) and stored at −80°C for future RNA purification.

### RNA purification and quality control

Total RNA was isolated using TRIzol Reagent (Thermo Fisher Scientific; cat. no. 15596018) and 1-bromo-3-chloropropane (BCP) (Molecular Research Center, cat. no. BP 151) phase separation method, followed by purification using the RNA Clean & Concentrator-5 kit columns (Zymo Research; cat. no. R1016). RNA quantity was measured with the RNA BR quantitation assays (Thermo Fisher Scientific; cat. no. Q10211) on a Qubit 2.0 fluorometer (Thermo Fisher Scientific; cat. no. Q32866). RNA quality was assessed with the RNA 6000 Nano chips (Agilent; cat. no. 5067-1511) on an Agilent 2100 Bioanalyzer system (Agilent; cat. no. G2939A).

### RNA sequencing

Small-RNA sequencing libraries were prepared with the NEXTflex small RNA-seq kit v3 (Bioo Scientific, cat# NOVA-5132-06). Gel size selection was performed separately for each PCR amplified barcoded sample, by cutting out the 150 bp band followed by bead cleanup. Barcoded libraries were normalized and pooled. Single-read sequencing (50 bp) was performed on an Illumina HiSeq 3000 sequencer (Illumina; cat. no. SY-401-3001).

Total-RNA sequencing libraries were prepared with the TruSeq Stranded Total RNA with Ribo-Zero Gold kit (Illumina; cat. no. RS-122-2303). Indexed libraries were normalized and pooled. For biological replicate 1, paired-end sequencing (2 × 94 bp) was performed on an Illumina HiSeq 2000 sequencer (Illumina; cat. no. SY-401-1001). For biological replicates 2–4, paired-end sequencing (2 × 75 bp) was performed on an Illumina HiSeq 3000 sequencer (Illumina; cat. no. SY-401-3001).

### Long non-coding RNA data analysis

Illumina base call (BCL) files from total-RNA sequencing were converted to FASTQ format with bcl2fastq2 v2.20 (Illumina, Inc.). Adapter sequences were trimmed with Cutadapt v1.10 (Martin, 2011) in Python v2.7.9, with the following adapter sequences as input:

Read 1: 5’-AGATCGGAAGAGCACACGTCTGAACTCCAGTCAC-3’
Read 2: 5’-AGATCGGAAGAGCGTCGTGTAGGGAAAGAGTGT-3’.

Trimmed reads under 20 bases were discarded. Quality control of adapter-trimmed sequences was performed on FastQC (Andrews, 2010). Adapter-trimmed FASTQ files were aligned to the reference human genome assembly (GRCh38) with STAR v2.7.9 (Dobin et al., 2013). The transcript annotation (GTF) file was obtained from GENCODE, release 43 (Harrow et al., 2012).

The binary alignment (BAM) files were used as input for de novo transcript assembly with the program StringTie v2.2.1 (Pertea et al., 2015). The inferred transcripts for each sample were then merged into a non-redundant set of assembled transcripts. The user-selected prefix ATRNS0523 was used for de novo assembled transcripts.

To distinguish known from potential novel transcripts, the program GFFcompare v0.11.6 (Pertea and Pertea, 2020) was used to contrast the de novo annotated transcripts to those in the GENCODE 43 annotation. This revealed 3,905 novel (not previously annotated) transcripts in our data. These were further selected, leaving only novel transcripts that were intergenic, fully contained within reference introns, or having exonic overlap with a reference exon but on the opposite strand (gffcompare categories u, i, or x, respectively). This final set of candidate novel transcripts was then merged with the GENCODE 43 annotation, yielding a reference GTF file of known (previously annotated) or novel (not previously annotated) transcripts.

The alignment (BAM) files were then used to generate a matrix of un-normalized read counts with the featureCounts program of the package Subread v1.5.3 (Liao et al., 2014). Paired-end exonic fragments were grouped at the level of genes, based on the reference GTF file of known or novel transcripts described above.

Differential expression analysis was performed with the R package DESeq2 v1.40.2 (Love et al., 2014). For each cell type and time point, differential expression analysis was performed contrasting GC-treated versus vehicle-treated cells, with the vehicle as the reference condition.

### Micro-RNA data analysis

Illumina BCL files from small-RNA sequencing were converted to FASTQ format using bcl2fastq2 v2.20 (Illumina, Inc.). Adapter trimming was first performed with Cutadapt v1.10 (Martin, 2011) in Python v2.7.9, with the following adapter sequence as input:

5’-TGGAATTCTCGGGTGCCAAGGAACTCCAGTCAC-3’

Trimmed reads less than 18 bases were discarded. The 5’ and 3’ four-nucleotide barcodes introduced by the library preparation chemistry were then removed with the Fastx Toolkit v0.0.14.

Quantification of known and novel mature microRNAs (miRNAs) was performed using the OpenOnmics mir-seek v0.3.0 pipeline (Kuhn and Zhang, 2024). This pipeline is composed of a series of data-processing and quality-control steps optimized for processing miRNA-sequencing data. As part of this pipeline, a second round of adapter-sequence trimming was performed with fastp v0.23.4 (Chen et al., 2018), using the minimum length option to remove reads with a length less than or equal to 16 bases. Quality control was performed on FastQC (Andrews, 2010) before and after trimming adapter sequences.

miRNA expression was quantified using the miRDeep2 suite v0.1.3 (Friedlander et al., 2012). To quantify known miRNA expression, the mapper.pl program was used to collapse and align reads against the human reference genome (GRCh38). Next, the miRDeep2.pl program was run to estimate the expression of known mature miRNAs using the human (hsa) v22 annotation from miRBase (Kozomara et al., 2019).

To quantify novel miRNA expression, miRDeep2 was run using a two-pass, cohort-level approach. In the first pass, the trimmed reads from all samples were concatenated into a single FASTA file. This cohort-level FASTA file was then passed to the mapper.pl program, to collapse and align the reads against GRCh38. The resulting alignment was provided as input to miRDeep2.pl to generate a cohort-level miRNA transcriptome FASTA file containing predicted novel mature miRNAs. In the second pass, this cohort-level miRNA FASTA file was provided to the quantifier.pl program to estimate the expression of each novel mature miRNA in each sample. The known and novel counts of each sample were then aggregated into two count matrices for downstream analysis.

The first matrix had 2,656 known miRNAs with raw read counts across the samples, of which 1,729 had at least one read in at least one sample. The second matrix had 1,994 candidate novel miRNAs with raw read counts across the samples, of which all had at least one read in at least one sample.

Differential expression analysis was performed with the R package DESeq2 v1.40.2 (Love et al., 2014). For each cell type and time point, differential expression analysis was performed contrasting GC-treated versus vehicle-treated cells, with the vehicle as the reference condition.

## Interactive figure

The interactive figure for exploring the results of this study was generated with the R package shiny (Chang et al., 2025).

## Data availability

The total RNA-seq and small-RNA-seq datasets will be publicly available through the Sequence Read Archive (SRA) and Gene Expression Omnibus (GEO) upon publication.

## Acknowledgements

This work was supported by the Intramural Research Program of the National Institute of Arthritis and Musculoskeletal and Skin Diseases, and by the Division of Intramural Research of the National Institute of Allergy and Infectious Diseases, both at the U.S. National Institutes of Health. This work utilized the computational resources of the NIH high-performance computing Biowulf cluster (https://hpc.nih.gov). We thank Dr. Justin Lack of the Integrated Data Sciences Section at the National Institute of Allergy and Infectious Diseases, National Institutes of Health, for coordination of bioinformatics services.

## NIH disclaimer

This research was supported by the Intramural Research Program of the National Institutes of Health (NIH). The contributions of the NIH authors are considered Works of the United States Government. The findings and conclusions presented are those of the authors and do not necessarily reflect the views of the NIH or the U.S. Department of Health and Human Services.

